# Testing Reversibility of Endosymbiotic Gene Transfer between Chloroplast and Nucleus

**DOI:** 10.64898/2026.07.03.736199

**Authors:** David Su, Sheng-An Chen, Philip Hammer, Eleena Chacko, Vadim Beilinson, Alexander Kinev, Masayuki Onishi

## Abstract

Most proteins targeted to the organelles of endosymbiotic origin are encoded in the nuclear genome, placing them under the regulatory dominance of the nucleus. For photosynthetic eukaryotes, nuclear-encoded chloroplast proteins arise via two routes: First, genes of cyanobacterial origin were relocated to the nucleus through endosymbiotic gene transfer (EGT). Second, proteins of eukaryotic origin emerged to support chloroplast function and structure. These proteins are reimported into the chloroplast via an import machinery. Reversing the transfer of such genes from the nucleus to the chloroplast genome may offer insights into chloroplast regulation and evolution. In this study, we established a highly efficient and accessible electroporation protocol for chloroplast transformation in the green alga *Chlamydomonas reinhardtii*, and used it to reverse-transfer two nuclear-encoded genes encoding proteins arising via the two routes described above: the cyanobacteria-derived chloroplast division protein FtsZ1 and the Rubisco-linker EPYC1 of eukaryotic origin. Regardless of origin, both chloroplast-encoded FtsZ1 and EPYC1 showed proper localization and functionality comparable to their nuclear-encoded counterparts. Together, our study provides a robust protocol for chloroplast transformation, a platform for investigating the evolutionary drivers of EGT, and a foundation for advancing chloroplast bioengineering.

**SIGNIFICANCE STATEMENT:**

- Endosymbiotic gene transfer has resulted in the mass migration of genes from the chloroplast genome to the nuclear genome. Reversing the gene transfer could reveal the evolutionary significance of genome partitioning.
- Using the green alga *Chlamydomonas reinhardtii*, this study developed an efficient, electroporation-based protocol for chloroplast transformation. Relocating the genes encoding two chloroplast-targeted proteins, *FTSZ1* and *EPYC1*, to the chloroplast genome showed that the proteins maintained normal localization and function.
- The established transformation protocol facilitates systematic testing of reverse gene transfer to elucidate the potential evolutionary advantages of genome partitioning and opens new avenues for chloroplast bioengineering.

## INTRODUCTION

Modern eukaryotic cells are products of ancient endosymbiotic events, in which once free-living prokaryotes were integrated into the cell as specialized, regulated organelles. For example, mitochondria derive from an ancestral alpha-proteobacterium that formed an endosymbiotic relationship with an archaeon, and plastids such as chloroplasts originated from a cyanobacterium that was acquired by an early eukaryote as an endosymbiont (Gould *et al*., 2008; Roger *et al*., 2017) (**Figure 1A, *Left***). While chloroplasts have retained the bulk of their prokaryotic biochemistry (Osteryoung and Vierling, 1995; Sharma *et al*., 2007), they have offloaded most of their ancestral bacterial genes to the host nucleus via endosymbiotic gene transfer (EGT; **Figure 1A, *Middle***) (Timmis *et al*., 2004). EGT presents significant evolutionary challenges and opportunities, as the physical escape and nuclear integration of the bacterial genes do not immediately provide functionality. To be successfully expressed, the newly integrated sequences must acquire eukaryotic regulatory elements, including nuclear promoters and polyadenylation signals, as well as N-terminal chloroplast-transit peptides (cTPs) (Martin and Herrmann, 1998). During this transition, these genes often exist in a state of redundancy in both the organelle and nuclear genomes (Adams *et al*., 1999; Choi *et al*., 2006); as the nuclear copy is successfully expressed, the selection pressure to retain the original copy disappears. In the modern land plants and green algae, only ∼120-150 genes are retained in the chloroplast genome (Green, 2011), which is less than 4% of the number of genes found in the widely studied cyanobacterium *Synechocystis* sp. PCC 6803 (Kaneko *et al*., 1996).

**FIGURE 1:**
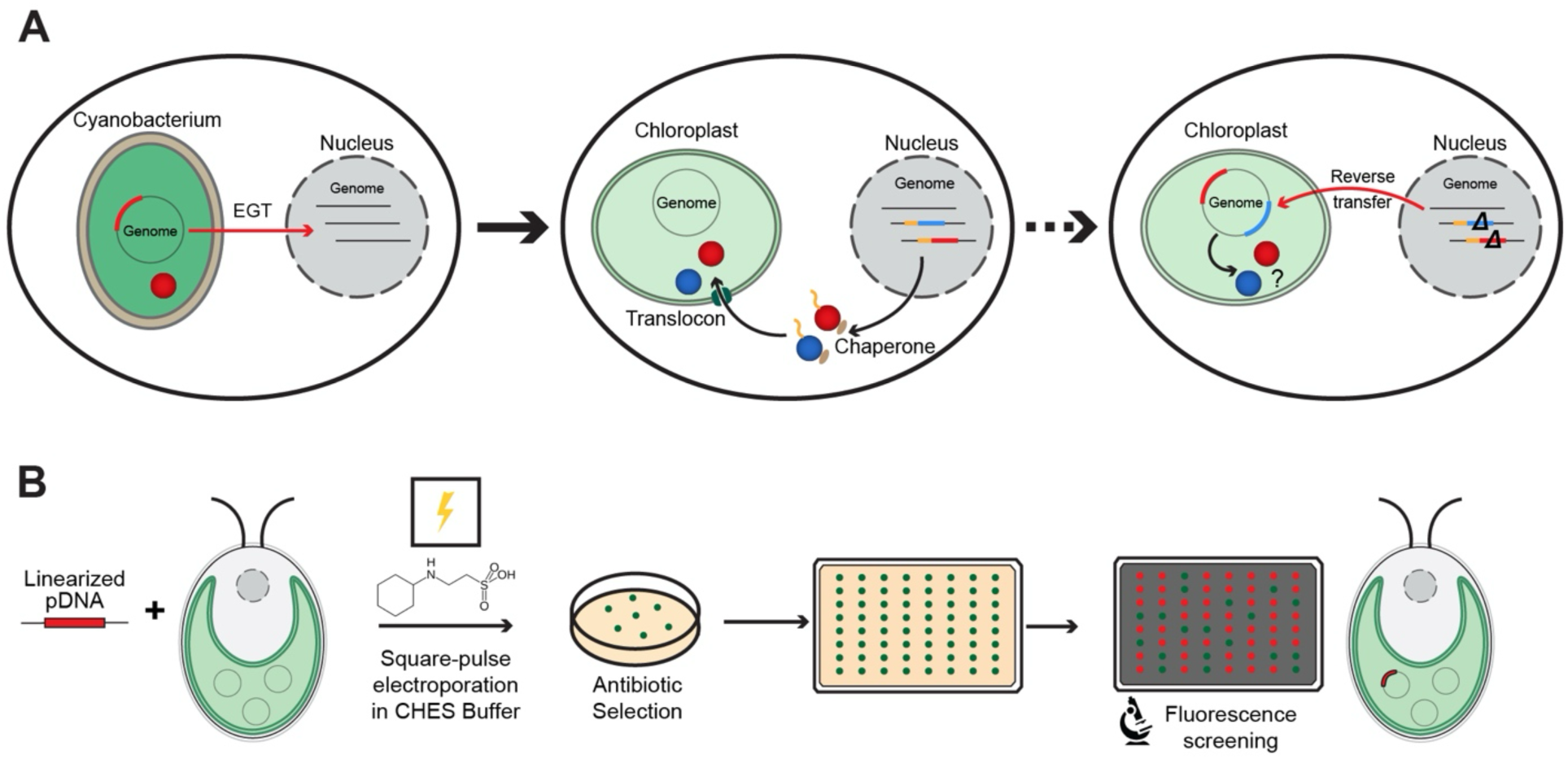
Reversing endosymbiotic gene transfer by chloroplast genome editing. (A) Throughout evolution, genes from the cyanobacterial ancestor of chloroplasts (red boxes) were transferred to the nuclear genome of the eukaryotic host (left panel). As a result, many chloroplast-targeted proteins (red circles) are synthesized in the cytosol and imported into the chloroplast through the translocons in the outer and inner chloroplast envelopes (middle panel), which is guided by a chloroplast transit peptide (yellow) and facilitated by cytosolic chaperones (brown). In addition, new genes encoding chloroplast-targeted proteins (blue boxes and circles) evolved in the eukaryotic host and are directed to the chloroplast via the same import route. In this study, we transferred nuclear-encoded chloroplast proteins into the chloroplast genome to examine whether the site of expression affects their functionalities and regulations (right panel). (B) An overview of the workflow for chloroplast transformation via electroporation in *Chlamydomonas reinhardtii*. Linearized plasmid DNA containing a chloroplast expression cassette (e.g., a fusion protein with a fluorescent marker) is transformed into *Chlamydomonas* with CHES buffer by square-pulse electroporation. Transformants are selected on antibiotic-containing medium, and resulting colonies are isolated and screened in a high-throughput manner using fluorescence microscopy. Positive clones are subsequently verified by genotyping, and the localization and functionality of the transgenic protein are examined.

After EGT, genes encoding chloroplast-targeted proteins are transcribed in the nucleus, translated into preproteins in the cytosol while kept fully or partially unfolded by cytosolic chaperones, and targeted for import through the chloroplast translocon complexes that span the outer and inner membranes (**Figure 1A, *Middle***) (Richardson and Schnell, 2020). Translocated preproteins are then processed to remove cTPs and folded by stromal chaperones, sometimes forming a complex with proteins expressed from the chloroplast genome. This modern configuration raises a fundamental question: What evolutionary advantages favor the segregation of genes into two separate compartments, despite the added complexity and associated energetic cost? Comparative genomics studies suggest that relocating organellar genes to the nucleus provides several theoretical evolutionary advantages, such as reduced mutation rate (Allen and Raven, 1996; Lynch, 1997), cellular control of chloroplast function (Woodson and Chory, 2008), and evolution flexibility through meiotic recombination (Martin and Herrmann, 1998). Reverse transfer of nuclear-encoded genes into chloroplast genomes has been used as an experimental approach to elucidate the driving force behind some specific cases of EGT (**Figure 1A, *Right***) (Whitney and Andrews, 2001; Dhingra *et al*., 2004). However, large-scale reverse gene transfer (RGT) is impractical, as traditional methods for DNA delivery into chloroplasts, such as biolistic transformation (Boynton *et al*., 1988; Narra *et al*., 2025), the PEG-protoplast method (O’Neill *et al*., 1993), and the glass-bead method (Kindle *et al*., 1991), are labor-intensive, cost-inefficient, and/or low in expression efficiency. Overcoming these barriers by developing a more accessible and efficient transformation platform is crucial for studying EGT. Such a method will also have translational value in bioengineering, as chloroplasts have been a target of plant and algal biotechnology (Bock, 2015; Kwon *et al*., 2018).

In this study, we developed an optimized chloroplast transformation platform for the model alga *Chlamydomonas reinhardtii,* using electroporation with a specialized DNA-delivery buffer as a highly efficient alternative for chloroplast gene delivery. We tested the feasibility of RGT by expressing two nuclear-encoded, chloroplast-targeted proteins of distinct evolutionary origins from the chloroplast genome: the bacterial-derived division protein FtsZ1 (TerBush *et al*., 2013) and the eukaryotic-evolved pyrenoid-forming protein EPYC1 (Mackinder *et al*., 2016). Surprisingly, chloroplast-encoded versions of both proteins localized correctly and appeared functionally equivalent to their nuclear-encoded counterparts. These results establish RGT as a powerful tool for studying EGT. Furthermore, the established platform may offer a robust chloroplast transformation method for bioengineering in photosynthetic eukaryotes.

## RESULTS

### Efficient chloroplast transformation by square-pulse electroporation in CHES buffer

To overcome the cost and technical limitations of traditional chloroplast transformation methods, we aimed to develop an efficient electroporation protocol for chloroplast transformation in *Chlamydomonas* (**Figure 1B)**. In short, plasmid DNA containing a spectinomycin-resistance cassette (Jackson *et al*., 2022) is linearized and mixed with cells in a CHES (N-cyclohexyl-2-aminoethanesulfonic acid)-based transformation buffer. The cells are then electroporated using a NEPA21 square-pulse electroporator, and the transformants are selected on spectinomycin-containing agar plates until colonies form. Using a reporter plasmid harboring a separate cassette encoding chloroplast-codon-optimized *mNEONGREEN* (*cpNG*), we quantified the efficiency of this method. After random colonies were picked, they were transferred to agar plates supplemented with amido black (which suppresses autofluorescence from media; Gutiérrez et al., 2022) and scanned with a wide-field fluorescence microscope equipped with a 5x long-working-distance objective. This electroporation protocol produced a high number of colonies with >75% expression rates (**Table 1**), significantly more than those of typical chloroplast transformation methods such as biolistic transformation. The CHES buffer system was previously developed to achieve efficient nuclear transformation with robust transgene integration and expression during electroporation of *Chlamydomonas* (Onishi and Pringle, 2016); here, it has also proven effective for DNA delivery into the chloroplast.

**Table 1.** Summary of chloroplast transformation efficiency by electroporation.

| cpNG plasmid<br>(pSC060, 2 $\mu\text{g}$) | Round | Spectinomycin-resistant colonies | NG-positive<br>colonies | Expression<br>rate |
| --- | --- | --- | --- | --- |
| Linearized (AflIII and StuI) | 1 | 54 | 41 | 76% |
| Linearized (AflIII and StuI) | 2 | 52 | 42 | 81% |
| Circular | 1 | 11 | 7 | 63% |
| Circular | 2 | 13 | 8 | 62% |

The chloroplast genome (plastome) consists of multiple identical circular nucleoid DNA copies (Kuroiwa *et al*., 1981; Gallaher *et al*., 2018). After CHES-mediated electroporation and spectinomycin selection, many colonies contained cells with heterogeneous cpNG signal levels (**Figure 2A, Passage 0**), suggesting a heteroplasmic plastome state in which transformed and untransformed nucleoids are present within the same chloroplast. Continued antibiotic selection and passaging are required to eliminate wild-type nucleoids and drive transformants toward homoplasmy. After serial restreaking under selective conditions at 24-hour intervals, the percentage of cells with detectable cpNG gradually increased from the initial 55% (Passage 0) to 75%, 92%, and 100% at Passages 1, 2, and 3, respectively (**Figure 2A**). PCR genotyping confirmed this trend: both the native chloroplast locus (“wild-type” band) and the cpNG-containing locus (“integration” band) were detected at early passages, while the wild-type band disappeared over time and the integration band increased in intensity (**Figure 2B**). To test the versatility of this protocol, we generated an independent strain expressing an additional fluorescent reporter, chloroplast-encoded *mSCARLET* (*cpSC*). After three passages, cpSC fluorescence was detectable in all transformant cells (**Figure 2C**). Efficient and rapid co-expression of cpNG and cpSC from an operon-like bicistronic construct using the *apcB*–*apcA* intercistronic element from the cyanobacterium *Synechococcus elongatus*, which has previously been used in combination with biolistic bombardment (Yeon *et al*., 2023), was also achieved, with 57% of colonies showing mNG fluorescence and 22% showing mSC after three passages (**Figure 2D**). Reaching homoplasmy after three passages/days is significantly faster than with traditional chloroplast transformation methods, which typically require several weeks (Guzmán-Zapata *et al*., 2016; Jackson *et al*., 2022). Together with the high initial positive expression rate (**Table 1**), these results suggest that electroporation with the CHES buffer efficiently delivers many DNA molecules into each chloroplast, thereby facilitating their integration into multiple nucleoid copies.

**FIGURE 2:**
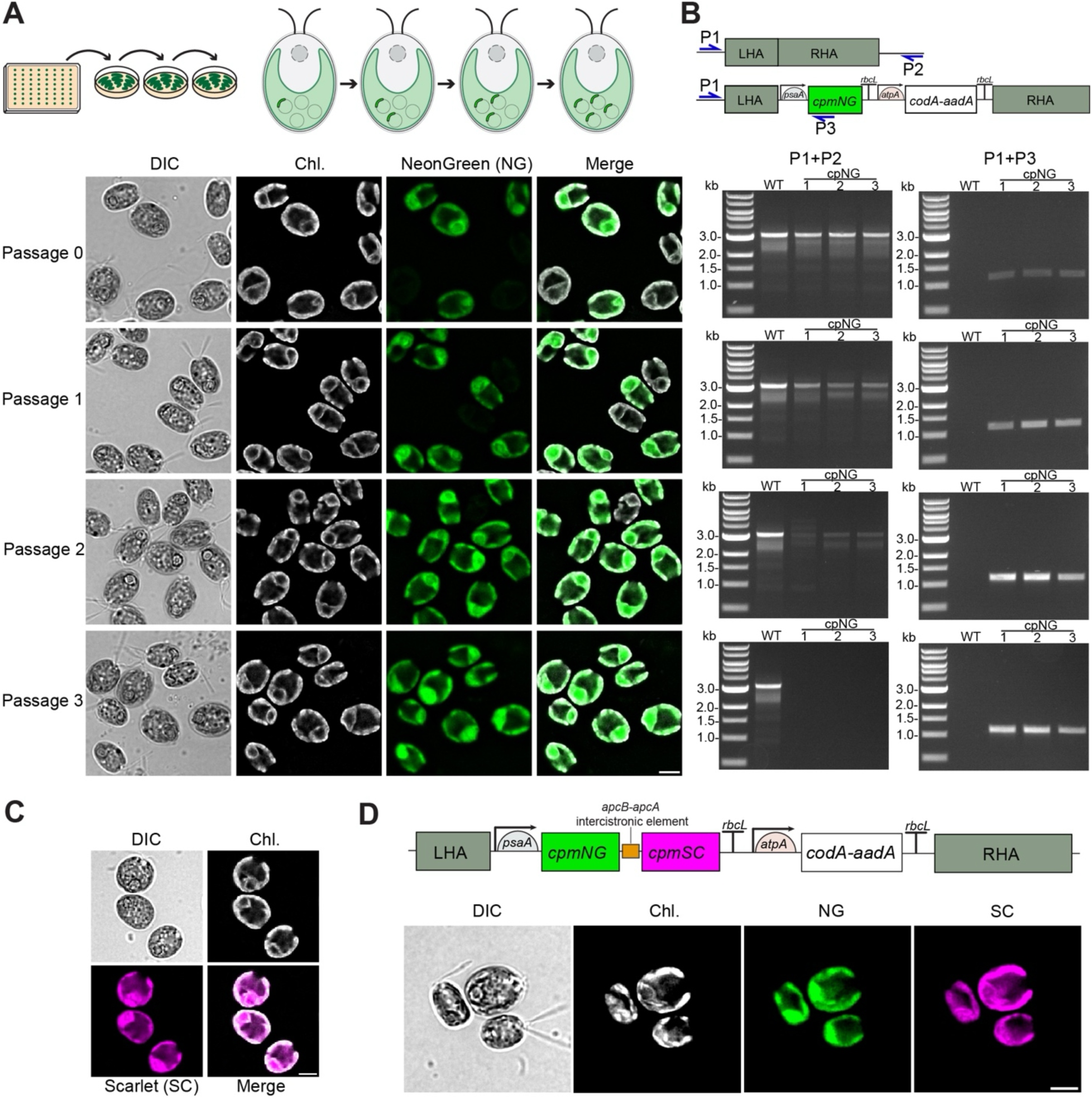
Successful delivery and expression of transgenic constructs via electroporation into *Chlamydomonas* chloroplast. (A) Fluorescence microscopy of cells expressing mNeonGreen from the chloroplast genome (cpNG) following square-pulse electroporation in CHES buffer. Differential interference contrast (DIC), chlorophyll autofluorescence (Chl.), mNeonGreen fluorescence (NG), and merged channels are shown. Images show serial passages (P0–P3), where passage 0 represents initial spectinomycin-resistant colonies recovered after electroporation and passages 1–3 represent successive restreaks under continuous selection at 24-hr intervals. Scale bar, 5 µm. (B) PCR genotyping of chloroplast transformants across passages showing amplification of the wild-type locus and integration-specific product. Early passages showed both products, indicating a mixed plastome population. Continued passaging led to loss of the wild-type band and enrichment of the integration product, consistent with progressive enrichment toward homoplasmy. (C) Fluorescence microscopy of an mScarlet reporter expressed from the chloroplast genome (cpSC) using the same electroporation-based transformation strategy. DIC, chlorophyll autofluorescence, cpSC fluorescence, and merged channels are shown. Scale bar, 5 µm. (D) Bicistronic expression of cpNG and cpSC from a single transcriptional unit. DIC, chlorophyll autofluorescence, cpNG fluorescence, and cpSC fluorescence are shown. Scale bar, 5 µm.

The design of our transgenic constructs is based on the CpPosNeg study (Jackson *et al*., 2022), in which circular plasmid DNA was directly transformed to integrate into the chloroplast genome via homologous recombination. Because bacterial and cyanobacterial genome editing by homologous recombination typically use linearized constructs to target specific loci (Golden *et al*., 1987; Datsenko and Wanner, 2000), we compared the transformation efficiency of cpNG plasmids that were either linearized by restriction digestion (as above) or remained circular. We observed that linearized DNA consistently yielded a higher number of colonies overall and a greater number of successful transgene expression per µg of DNA used (**Table 1**). Given the marked increase in efficiency, linearization was adopted for all subsequent experiments in this study.

### Chloroplast-expressed FtsZ1 assembles into a division ring of normal morphology and dynamics

We next tested the feasibility of RGT by transferring a nuclear-encoded gene encoding a chloroplast-targeted protein into the plastome. We focused on FtsZ1, a protein of cyanobacterial origin that is a key component of the chloroplast division apparatus (Osteryoung *et al*., 1998; Schmitz *et al*., 2009; TerBush *et al*., 2013). Like their bacterial ancestors, chloroplasts divide through binary fission driven by the constriction of the stromal FtsZ ring at the midzone. FtsZ is a cytoskeletal GTPase with structural similarity to tubulins (Erickson, 1997); in green algae and land plants, paralogs FtsZ1 and FtsZ2 co-assemble into heteropolymeric protofilaments to form the FtsZ ring (Miyagishima *et al*., 2004; TerBush *et al*., 2013) (**Figure 3A**). As part of EGT, the genes encoding components of the chloroplast division apparatus have been transferred to the nuclear genome in many species, including *FTSZ1* and *FTSZ2 in Chlamydomonas* (Miyagishima *et al*., 2012).

**FIGURE 3:**
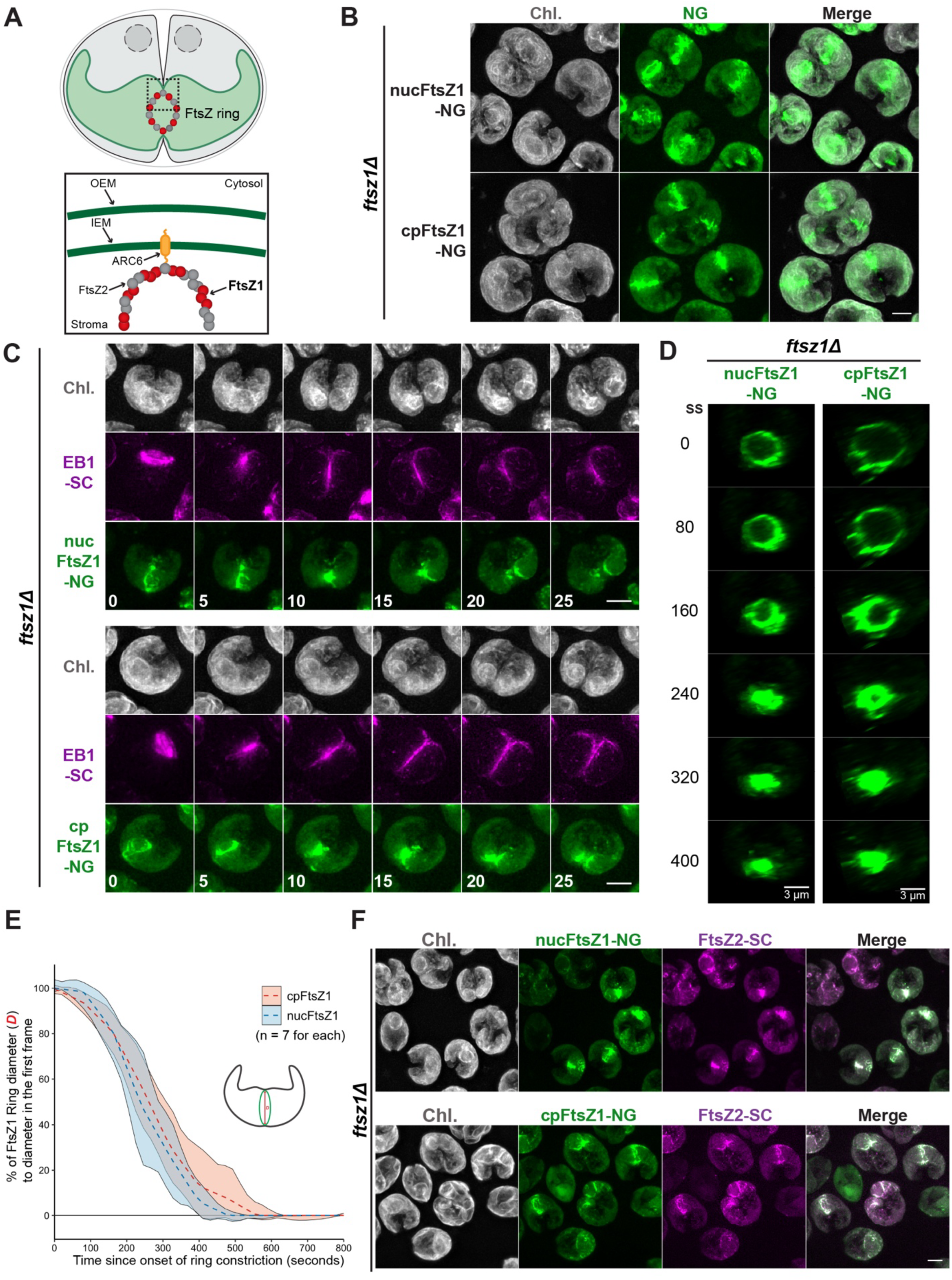
FtsZ1 forms a division ring of normal morphology and dynamics when expressed from the chloroplast genome. (A) Schematic of a dividing chloroplast in a *Chlamydomonas* cell, showing the FtsZ ring at the division site. The magnified view shows the division machinery in the chloroplast stroma. FtsZ1 (red) and FtsZ2 (gray) assemble to form the FtsZ ring, which is anchored to the chloroplast inner envelope membrane (IEM) through ARC6 (yellow). OEM, outer envelope membrane. (B) Images of *ftsz1Δ* cells expressing either nuclear (nuc)- or chloroplast (cp)-encoded FtsZ1-NG. Scale bar, 5 µm. (C) Time-lapse images of *ftsz1Δ* cells expressing either nuclear- or chloroplast-encoded FtsZ1-NG during cell and chloroplast division. Time is in minutes and set to zero when the mitotic spindle is observed, as shown by EB1-Scarlet. Scale bars, 5 µm. The full series are presented in Supplemental Videos S1 and S2. (D) Time-lapse comparison of constriction dynamics between nucFtsZ1-NG and cpFtsZ1-NG rings. Rotated 3D projections provide a face-on view of the FtsZ1 rings over time. Time in seconds (ss). Scale bars, 3 µm. The full series are presented in Supplemental Videos S3 and S4. (E) Comparison of the kinetics between nucFtsZ1-NG and cpFtsZ1-NG ring constrictions. FtsZ1 ring diameter (µm) at each given time point was measured and expressed as a percentage relative to the first frame (metaphase) over time. Means and 95% CIs of 1000×-bootstrapped locally estimated scatterplot smoothing (LOESS) curves are shown. (F) Images of *ftsz1Δ nucFTSZ2-SC* cells expressing either nucFtsZ1-NG or cpFtsZ1-NG. Scale bar, 5 µm.

We previously developed a live-cell imaging method for fluorescently tagged markers of cell and chloroplast division in *Chlamydomonas* (Clark-Cotton and Chen, *et al*., 2025), including FtsZ1-NeonGreen (NG). The expression construct has *FTSZ1’s* own promoter and is integrated into the nuclear genome; thus, we will hereby refer to the fusion protein as nucFtsZ1-NG. nucFtsZ1-NG is expressed in late-G1/S phase, similarly to endogenous FtsZ1 (Tulin and Cross, 2015; Zones *et al*., 2015; Strenkert *et al*., 2019), imported into the chloroplast stroma, and forms a ring at the chloroplast midzone in preprophase, both in wild-type background (Clark-Cotton and Chen *et al*., 2025) and in an *ftsz1Δ* mutant generated by CRISPR-Cas9-mediated knockout (**Figure 3B**). When FtsZ1-NG was expressed from the chloroplast genome (cpFtsZ1-NG) of *ftsz1Δ* cells, it assembled into a ring in a timely manner, showing no obvious differences from nucFtsZ1-NG **(Figure 3B**). cpFtsZ1-NG is transcribed by the constitutively active *psaA* promoter, and therefore its signal accumulates in interphase cells; however, cpFtsZ1-NG remained dispersed throughout the chloroplast stroma (**Supplemental Figure S1**), suggesting that other components of the ring encoded in the nuclear genome (e.g., FtsZ2 and ARC6) are required for Z-ring assembly, or that cell-cycle-regulated signaling triggers Z-ring formation, or both.

Next, we compared the dynamics of Z-ring constriction between cells expressing nucFtsZ1-NG and cpFtsZ1-NG. The timing of mitosis and cytokinesis was monitored using mScarlet-tagged EB1, a microtubule plus-end-binding protein (Onishi *et al*., 2020; Pecani *et al*., 2022; Clark-Cotton and Chen *et al*., 2025). Both chloroplast- and nuclear-encoded FtsZ1-NG formed a ring at the chloroplast midzone shortly before mitotic-spindle formation; the ring constricted as the furrow ingressed (**Figure 3C; Supplemental Videos 1 and 2**). Time-lapse imaging with short intervals showed no observable differences between the two types of FtsZ1 rings in the cells examined (**Figure 3, D and E; Supplemental Videos 3 and 4**). These results indicate that FtsZ1-NG is successfully integrated into the Z-ring, in the absence of endogenous FtsZ1, regardless of the expression origin.

The process of EGT likely involved a transitional phase in which genes encoding some components of a molecular apparatus or biological pathway were partially relocated to the nucleus, while others remained in the chloroplast genome. To determine whether FtsZ ring components can still co-assemble when partitioned into different genomes, we co-expressed cpFtsZ1-NG and nucFtsZ2-SC in the *ftsz1Δ* mutant. The cpFtsZ1 and nucFtsZ2 signals showed clear co-localization throughout different stages of chloroplast division (**Figure 3F**), suggesting the two proteins could still assemble into a heteropolymeric ring despite being expressed from distinct genomic compartments, thereby representing a plausible intermediate state of EGT. Taken together, these results suggest that FtsZ1 remains functionally intact when reverse-transferred to the chloroplast genome.

### Expression of EPYC1 from the chloroplast genome supports pyrenoid function

While FtsZ1 is a nuclear-encoded chloroplast protein of cyanobacterial origin, many other chloroplast-targeted proteins emerged in the nuclear genome after plastid acquisition to support newly evolved functions such as protein import (Richardson and Schnell, 2020) and stress response (Kodama and Sano, 2006). We next aimed to test whether such proteins of eukaryotic origin can be transferred to the chloroplast genome, effectively mimicking a de novo gene transfer in the reverse direction to canonical EGT. To this end, we targeted the pyrenoid-based carbon-concentration mechanism (CCM), another eukaryotic innovation driven by novel proteins that lack sequence homologs in cyanobacteria (He *et al*., 2023). The pyrenoid is a specialized, phase-separated compartment within the chloroplast that serves as the core of the CCM in algae and hornworts (Pritchard *et al*., 2026). In *Chlamydomonas*, a dedicated linker protein, EPYC1 (also known as LCI5), interacts multivalently with Rubisco, aggregating the enzyme into a dense matrix with concentrated CO₂ levels to enhance carbon fixation (Burow *et al*., 1996; Miura *et al*., 2004; Mackinder *et al*., 2016; **Figure 4A**); loss of EPYC1 disrupts Rubisco aggregation in the pyrenoid and impairs photoautotrophic growth (Mackinder *et al*., 2016).

**FIGURE 4:**
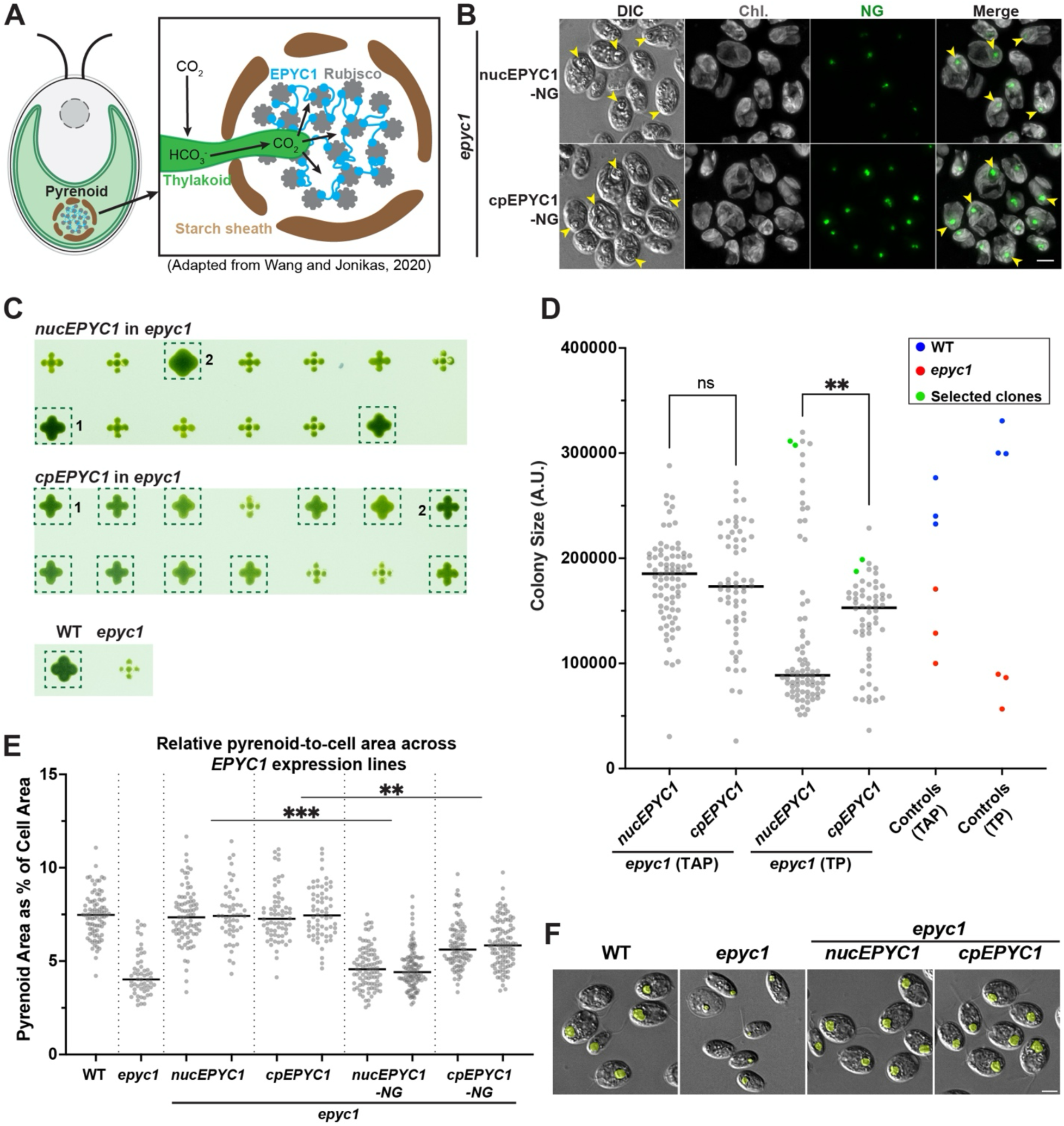
Expression of EPYC1 from the chloroplast genome supports pyrenoid function. (A) Schematic of the pyrenoid within the *Chlamydomonas* chloroplast. The magnified view shows the pyrenoid with EPYC1 (blue) linking Rubisco complexes (gray) to form a matrix. The pyrenoid is surrounded by a starch sheath and traversed by thylakoid tubules. CO₂ is converted to HCO_3_^-^, which is transported to the pyrenoid and converted back to CO₂ to support Rubisco-mediated carbon fixation. Adapted from Wang and Jonikas, 2020. (B) Localization of fluorescently tagged EPYC1-NG expressed from either the nuclear (nuc) or chloroplast (cp) genome in an *epyc1* mutant (CC-5360). Cells were grown on TP (tris-minimal) medium at low CO_2_ (ambient air). Arrowheads point to some of the pyrenoids in the chloroplasts. Both ncEPYC1-NG and cpEPYC1-NG showed comparable localization positioned inside the chlorophyll autofluorescence (Chl.). Scale bar, 5 µm. (C) Comparison of growth phenotypes between *nucEPYC1* and *cpEPYC1* transformants (without tag) in the *epyc1* mutant. The colonies were grown on TP (Tris-minimal) plates for 4 days at low CO_2_ (ambient air) under continuous light (300 μmol photons·m^−2^·s^−1^) before imaging. Colonies showing faster growth relative to *epyc1* are highlighted with dashed boxes. Numbers indicate the selected clones used in Fig. 4EF. WT, CC-5325. *epyc1,* CC-5360. (D) A scatter plot showing colony sizes of the *nucEPYC1* and *cpEPYC1* transformants cultured on TAP and TP plates. Each dot represents an individual transformant (n = 80 for *nucEPYC1* and n = 60 for *cpEPYC1*). Green dots indicate the selected clones used in Fig. 4EF. Blue and red dots represent three independent colonies of CC-5325 and CC-5360, respectively. The horizontal bars indicate the medians of each dataset. ns, not significant; **, p < 0.01 (Mann-Whitney test). (E) Comparison of pyrenoid sizes between WT, *epyc1*, and transgenic lines (two clones for each line) complemented with either *nucEPYC1* or *cpEPYC1*. The cells were grown on TP medium. The total pyrenoid area as a percentage of the total cell area is plotted; each dot represents a single cell. From left (WT) to right (*cpEPYC1-NG*), n = 86, 56, 84, 52, 64, 64, 93, 108, 92, 107. The horizontal bars indicate the medians of each dataset. WT: CC-5325. *epyc1*: CC-5360. **, p < 0.01; ***, p < 0.001 (Nested one-way ANOVA comparing expression constructs) (F) Representative images showing the pyrenoids in WT, *epyc1*, and *epyc1* complemented by *nucEPYC1* and *cpEPYC1.* The cells were grown on TP medium. The pyrenoids in the DIC images are highlighted in yellow. Scale bar, 5 µm.

To ask whether expressing EPYC1 from the chloroplast genome (cpEPYC1) affects its localization and pyrenoid formation, and therefore the carbon-fixing capacity of the transformed cells, we introduced the chloroplast-encoded *EPYC1* (untagged) and *EPYC1-NG* transgenes to an existing *epyc1* mutant (Mackinder *et al*., 2016). The cpEPYC1-NG signals formed a robust aggregate within the chloroplast and localized to the pyrenoid at the base of the cup-shaped chloroplast (as visualized by DIC microscopy; **Figure 4B**). During this experiment, we noticed that the sizes of pyrenoids induced by fluorescently tagged EPYC1, regardless of their expression origin (nucEPYC1-NG and cpEPYC1-NG), were significantly smaller than those supported by untagged counterparts (see **Figure 4, E and F**), suggesting that C-terminal tagging slightly compromises the protein function. Thus, we used untagged nucEPYC1 and cpEPYC1 in all subsequent experiments.

Next, we evaluated whether cpEPYC1 could restore defective growth in the *epyc1* mutant under photoautotrophic conditions, in which atmospheric CO₂ serves as the sole carbon source. We separately transformed the *epyc1* mutant with untagged *nucEPYC1* and *cpEPYC1* transgenes, isolated all the resulting colonies, and subjected them to photoautotrophic growth assays. As shown in Figure 4C for a subset of the colonies, only ∼30% of the *nucEPYC1* transformants showed restored growth on Tris-Phosphate (TP) medium without an external carbon source (**Figure 4C, top; Figure 4D**). Strikingly, the rate of restored growth was ∼80% for the *cpEPYC1* transformants (**Figure 4C, middle; Figure 4D**). Analysis of colony sizes for all isolated colonies (81 for *nucEPYC1* and 61 for *cpEPYC1*) as a proxy for growth showed that *cpEPYC1* supported higher overall growth compared to *nucEPYC1* (**Figure 4D**).

The *epyc1* mutant has a reduced pyrenoid size due to its inability to aggregate Rubisco into a pyrenoid matrix (Mackinder *et al*., 2016). To determine whether cpEPYC1 expression restores normal pyrenoid size in the *epyc1* mutant, we selected two independent clones from the subset of *cpEPYC1, nucEPYC1*, *nucEPYC1-mNG*, and *cpEPYC1-mNG* transformants that showed improved growth on TP medium and quantified their pyrenoid sizes. The results showed that *nucEPYC1* and *cpEPYC1* are equally capable of supporting pyrenoid formation (**Figure 4, E and F**). Interestingly, when mNeonGreen-tagged nucEPYC1 and cpEPYC1 were compared, although neither was fully functional, the latter formed slightly larger pyrenoids, perhaps due to stronger or more stable expression.

In summary, these results show that reverse-transferred cpEPYC1 supports pyrenoid formation and cell growth under photoautotrophic conditions. A parallel comparison between *nucEPYC1* and *cpEPYC1* transformants shows that chloroplast expression has higher complementation rates and phenotypic consistency, highlighting it as a powerful approach for introducing heterologous proteins to engineer organelle functions.

## DISCUSSION

### Efficient electroporation expands the toolkit for *Chlamydomonas* chloroplast bioengineering and potentially beyond

Chloroplasts have been targeted for engineering, serving as a biofactory to produce biopharmaceuticals, nutritional supplements, and other high-value biomolecules, as well as platforms for enhancing plant and algal traits (Kwon *et al*., 2018; Jackson *et al*., 2021; Narra *et al*., 2025). However, editing chloroplast genomes has traditionally relied on biolistic transformation (Narra *et al*., 2025), a method that requires specialized equipment and expensive consumables, thereby limiting widespread adoption and experimental throughput.

In this study, we showed that chloroplast transformation via CHES-mediated electroporation in *Chlamydomonas* resulted in a positive screening rate of over 70%, with the majority of drug-resistant transformants successfully expressing the transgenic fluorophore (**Table 1**). Furthermore, fluorescence analysis confirmed progressive enrichment toward homoplasmy within three serial passages under continued selection (**Figure 2A and 1B**). The accessibility of this protocol, together with high positive rates and rapid establishment of uniform transgene expression, facilitates high-throughput chloroplast engineering.

Although electroporation-mediated nuclear transformation has been used in multiple microalgal species with high efficiency and does not require cell wall removal (Gutiérrez and Lauersen, 2021), electroporation for chloroplast transformation has only been established in *Chlorella vulgaris* (Bolaños-Martínez *et al*., 2022). Our high-efficiency electroporation method should in principle be applicable to many other algal species with minor modifications.

### Advantages of direct chloroplast engineering

In bioengineering, plastid expression offers unique advantages over nuclear expression, regardless of the transgene’s origin (chloroplast, nuclear, or heterologous). First, a transgene can be integrated into specific loci through homologous recombination mediated by a bacterial-like RecA protein (Nakazato *et al*., 2003), thereby bypassing the unstable expression that can result from random nuclear integrations. The high transgene copy number in homoplasmic lines further supports robust expression. Second, the chloroplast’s prokaryotic heritage allows for polycistronic expression, which achieves coordinated synthesis of proteins within a biological pathway or subunits of a complex. Additionally, codon usage analyses among green eukaryotes demonstrate a shared preference for A- and T-ending codons (Nakamura *et al*., 1999; Qi *et al*., 2015); thus, a transgene optimized for one species is likely to be transferable to another. Third, the predominant maternal inheritance of chloroplast genomes in green algae and land plants serves as a natural biocontainment system, preventing accidental gene flow via paternal gametes (Schneider, 2023). Conversely, rare or induced events of biparental transmission, as observed in various species such as *Nicotiana tabacum* (Chung *et al*., 2023) and *Chlamydomonas* (Joo *et al*., 2022), could also be exploited (Yadav *et al*., 2025). Lastly, many algal species of environmental, industrial, medical, and academic importance [e.g., *Symbiodinium minutum* (Liu *et al*., 2018), *Coccomixas p.* (Kasai *et al*., 2018), *Nannochloropsis oceanica* (Vieler *et al*., 2012), *Chromochloris zofingiensis* (Gallaher and Roth, 2018), *Porphyridium purpureum* (Bhattacharya *et al*., 2013), *Porphyra umbilicalis* (Brawley *et al*., 2017), *Klebsormidium flaccidum* (Hori *et al*., 2014)] have limited genome engineering tools available. Direct editing of chloroplast genomes instead of the nuclear genome may provide a shortcut for modifying traits in these species.

### Chloroplast genome editing can shed light on its function and evolution

The physical migration of organelle genes into the nucleus through EGT is a ubiquitous, ongoing driver of eukaryotic genome evolution (Martin, 2003; Zhong *et al*., 2026). When we reversed the migration of *FTSZ1*, a gene of cyanobacterial origin, by expressing it from the chloroplast genome, the resulting transformants exhibited no obvious defects in chloroplast division or cell viability. The proper colocalization of cpFtsZ1 with the nuclear-encoded binding partner (FtsZ2) of the division apparatus and the normal dynamics of the FtsZ ring during constriction suggest that the chloroplast stroma can support various processes required for FtsZ1 function, such as its synthesis, folding, complex formation with FtsZ2, ring formation, and contraction. The data also suggest that the cell-imposed regulations, such as cell-cycle-regulated transcription, post-translational modification in the cytosol, and controlled translocation into the chloroplast, are not crucial for FtsZ1 function, at least under the experimental conditions used in this study. This raises an evolutionary question: Why was FtsZ1 transferred to the nuclear genome in the first place?

Equally surprising is that the expression of *cpEPYC1* is also well-tolerated by the cell. Unlike FtsZ1, EPYC1 has no cyanobacterial ancestry; it is a eukaryotic innovation that evolved within the nuclear genome to induce aggregation of Rubisco. This unexpected tolerance suggests that reverse transfer of genes from the nucleus to the chloroplast *can* occur and raises an evolutionary question: Why do many chloroplast proteins of non-cyanobacterial origin appear to have emerged only in the nucleus, been stably retained there, and never transferred to the chloroplast genome?

Ultimately, elucidating these questions about chloroplast evolution requires further RGT-type experiments (e.g., Whitney and Andrews, 2001) conducted systematically and at high throughput. Efficient chloroplast transformation by square-pulse electroporation using CHES buffer should facilitate such studies.

## MATERIALS AND METHODS

### Strains and growth conditions

For transformations and expression of cpNG, cpNG-SC, and cpFtsZ1-NG, *Chlamydomonas reinhardtii* wild-type strain iso10 (mt+, congenic to CC-124; provided by S. Dutcher, Washington University in St. Louis, St. Louis) was the parental strain. For transformation of EPYC1-NG, cpEPYC1, and cpEPYC1-mNG, *C. reinhardtii* strains CC-5325 (wild type) and CC-5360 (*epyc1*) (Mackinder *et al*., 2016) were the recipient strains. Routine cell culture was done in Tris-acetate-phosphate (TAP) medium at ∼25°C under constant illumination (50 μmol photons·m^−2^·s^−1^). Tris-phosphate (TP) medium with no external carbon source was used to compare the growth of transgenic *EPYC1* strains under photoautotrophic conditions. Genetic crosses were done as previously described (REF). All *C. reinhardtii* strains used in this study can be found in Supplemental Table 1.

### Plasmid constructions

All chloroplast expression plasmids used in this study are listed in Supplemental Figure S2 and Supplemental Table 2, and their sequences are available in Supplemental File 1. The plasmids were constructed using Start-Stop Assembly (Taylor *et al*., 2019) and/or Gibson assembly (Gibson, 2009). They share the basic backbone structure with the cpPosNeg plasmid pLuc2 (Jackson *et al*., 2022), with a *codA-aadA* fusion marker that can be selected and counter-selected with spectinomycin and 5-fluorocytocine, *psaA/rbcL* promoter/terminator for transgene expression, and left and right homology arms (LHA and RHA) for recombination at the *psbH* gene-downstream locus, except that RHA in our constructs is extended to ∼2.4 kb (∼1.2 kb in pLuc2). Inserts of interest, which were codon-optimized using SnapGene according to the *C. reinhardtii* chloroplast codon usage table (Kazusa Codon Usage Database) and synthesized by Twist Bioscience, include: *cpmNEONGREEN*, *cpmSCARLET*, *cpFTSZ1* (Cre02.g118600), and *cpEPYC1* (Cre10.g436550). For *cpFTSZ1* and *cpEPYC1*, the coding sequences corresponding to the mature proteins after cTP processing were used. The cTP cleavage site of FtsZ1 was predicted using TargetP 2.0 (Almagro Armenteros *et al*., 2019). The putative cleavage site of EPYC1’s cTP was based on the smallest mature proteoform previously described (Atkinson *et al.,* 2019).

For making the nucEPYC1-NG expression plasmid pDS023, the pLM022 plasmid (*P_psaD_:EPYC1-mCherry-6xHIS*) (Mackinder *et al*., 2016) was linearized by PCR-amplification without the *mCherry* sequence. The resulting backbone was joined with a *NG_v2-3FLAG* fragment (PCR-amplified from template plasmid pRT067) via Gibson assembly. For making the nucEPYC1 expression plasmid pDS024 without tag, the *mCherry* sequence was removed from pLM022 by PCR-amplification using two primers that bind at the end of *EPYC1* CDS (reverse) and after the stop codon of *mCherry-6xHIS* CDS (forward); the primers contain overlapping sequence that allows the linear backbone to self-circularize through Gibson assembly.

### Chloroplast transformation by CHES-mediated electroporation

Plasmids were linearized with StuI and AflII at two cut sites inside the *psbH* homology arms (Supplemental Figure 2). 2 µg of plasmid DNA was mixed with NEBuffer 4 (NEB, B7004S) and the enzymes in a final volume of 10 µL. *C. reinhardtii* cultures were grown in liquid TAP at 25°C under continuous light to ∼5 × 10^6^ cells/mL. The cells were collected by centrifugation, washed twice with CHES-EP buffer (10 mM *N*-cyclohexyl-2-aminoethanesulfonic acid, pH 9.25, 40 mM sucrose, and 10 mM sorbitol) and resuspended in the same buffer at 8 × 10^8^ cells/mL. For electroporation, 125 µL of cells was mixed with 10 µL of the digested plasmid; a 125 µL of the final mixture was then transferred to a 2-mm gap electrocuvette (Bulldog Bio, #12358-346), followed by electroporation using a NEPA21 square-pulse electroporator (Nepa Gene/Bulldog Bio. Poring pulses: 2 pulses, at 250 V and 150 V for 8 ms each. Transfer pulses: 5 pulses, starting at 20 V with a decay rate of 40% for 50 ms each. Expected energy is 1.5∼2 J). Immediately following electroporation, the cells were transferred into recovery medium (liquid TAP supplemented with 80 mM sucrose). The culture was incubated overnight at room temperature under dim light with continuous shaking, and the recovered cells were plated onto selection plates containing 100 µg/mL spectinomycin (Enzo Life Sciences, BML-A281-0010). Colonies that appeared after 7∼10 days were examined under fluorescence microscopy to assess successful expression of the construct. Positive colonies were then re-streaked onto new spectinomycin plates daily for 3 consecutive cycles to ensure that the plastome has reached homoplasmy.

### High-throughput fluorescence screening of transformants on amido black plates

Antibiotic-resistant colonies of chloroplast or nuclear transformations were picked using a Singer Instruments PIXL robot onto a TAP-agar plate containing appropriate antibiotics as a 96-array, allowed to recover for 3∼4 days, and then replicated (Singer Instruments ROTOR) onto a TAP-agar plate containing amido black (150 mg L-1), which reduces background autofluorescence (Gutiérrez et al., 2022). Cells on amido-black plates were allowed to grow for 12-24 hours and mounted to a Lecia Thunder inverted microscope and scanned in using an N PLAN 5X/0.12 PH0 objective and the following LED excitations and emission filters: 510 nm and 535/15 nm for NeonGreen (100%, 1 second), 550 nm and 595/40 nm for Scarlet (100%, 1 second); 640 nm and 705/72 nm for chlorophyll autofluorescence (100%, 20 ms).

### CRISPR-Cas9 mediated knockout of *FTSZ1*

Single guide RNA targeting the *FTSZ1* (Cre02.g118600) coding sequence (5’-GGGTAGGAGGTGCTCGTCGCGGG-3’) was synthesized by IDT (Integrated DNA Technologies) and dissolved in TE buffer at 100 µM. For the RNP complexing reaction, 1.2 µL of sgRNA was mixed with 2.8 µL of IDT Duplex buffer and 1.0 µL of 10 mg/mL Cas9 protein (Alt-R Cas9v3, IDT #1081058), and incubated at 37°C for 30 minutes. All subsequent procedures were as described previously (Wang *et al*., 2023).

### Light microscopy

Fluorescence microscopy images were collected on a Leica Thunder inverted microscope equipped with an HC PL APO 63X/1.40 N.A. oil-immersion objective lens. Fluorescence signals were captured using the following combinations of LED excitation and emission filters: 510 nm and 535/15 nm for NeonGreen; 550 nm and 595/40 nm for Scarlet; 640 nm and 705/72 nm for chlorophyll autofluorescence, with 0.4 μm Z-spacing covering 8 μm. All images were processed through Thunder Large Volume Computational Clearing and Deconvolution (Leica). Fluorescence images were converted to maximum projections using ImageJ (https://imagej.net/software/fiji/). DIC images were also captured on Leica Thunder and the medial plane of DIC Z-stack was selected to produce representative single-plane images. For time-lapse imaging of live cells expressing nucFtsZ1-NG or cpFtsZ1-NG, the cells were first synchronized with alternating 12-hour light: 12-hour dark (12L:12D) cycles (250 μmol photons·m^−2^·s^−1^) on TAP-agar plates for 2 days. At hours 10.5-11 of the 12L:12D cycle, the cells on the TAP-agar were collected and placed onto a 1.5% low-melting agarose block in TAP (IBI Scientific #IB70051). The agarose blocks were placed into wells of a chambered glass coverslip (Ibidi, 81817), and the space between the agarose block and the edge of the well was sealed with 2% low-melting agarose. The time-lapse experiment started at hours 11.5-12, and images were captured at 5-minute intervals. For time-lapse imaging of FtsZ1-NG ring constriction, images were captured at 80-second intervals for 45 minutes; Z-stack images were reconstructed into 3D using Imaris software (v10.2.0, Oxford Instruments). To track the constriction dynamics, the FtsZ1 ring diameter of a given cell was determined at each time point by measuring the distance (µm) between the two outermost poles of the FtsZ1-NG signal overlapping with the chlorophyll autofluorescence at the chloroplast division site (“Measurement Points” function of Imaris). The time point at which the mitotic spindle forms (EB1-mScarlet) was set as time zero.

### Colony-size quantification under photoautotrophic growth

For the colony growth assays, *nucEPYC1* and *cpEPYC1* transformants were picked onto a TAP-agar plate as a 96-colony array using a PIXL colony-picking robot (Singer Instruments), and subsequently replicated onto fresh TP- and TAP-agar plates using a ROTOR robot (Singer Instruments). For colony-size quantification, unprocessed RAW images of the TP-agar plates were captured using a digital camera at a fixed distance after four days of growth (one day for TAP-agar) under continuous light (300 μmol photons·m^−2^·s^−1^) in low CO_2_ (∼0.04% in ambient air). Captured RAW files were converted to RGB TIFF images. Colony quantification was processed using a custom ImageJ macro script: regions of interest (ROIs) for the 96-colony array were generated using manually aligned grids. The channels of the captured RGB images were split, and the raw integrated density of the green channel within each ROI was measured as a quantitative proxy for colony growth.

### Pyrenoid size quantification

All image processing and quantification were performed in Fiji/ImageJ. The DIC channel was used to determine whole-cell boundaries: the medial plane was thresholded using Huang’s fuzzy thresholding method (Huang and Wang, 1995), which emphasizes overall object boundaries rather than internal signal structure. This approach enabled robust detection of whole-cell outlines. Individual whole-cell masks were then generated using *Analyze Particles* in ImageJ, and the area of each cell was measured using the *Measure* function. Pyrenoids were identified in the DIC medial plane by the boundary formed by the surrounding starch sheath. Pyrenoid localization was confirmed using the chlorophyll autofluorescence channel, where the pyrenoid region typically appears as a localized reduction in autofluorescence signal relative to the surrounding chloroplast. Cells containing more than one pyrenoid or lacking a detectable pyrenoid were excluded from analysis. For each cell, the pyrenoid region was manually outlined, and the pyrenoid area was measured using the *Measure* function. The size of the pyrenoid in each cell was expressed as the ratio of the pyrenoid area to the area of the cell.

## ACKNOWLEDGMENTS

We thank Luke Mackinder for providing plasmid pLM022, Ryutaro Tokutsu for sharing the *mNEONGREEN_v2* sequence, Lisa Cameron and the Duke Light Microscopy Core Facility for their support and assistance with image analysis, and Matt Laudon and the Chlamydomonas Resource Center (https://www.chlamycollection.org/) for strains and plasmids. This center is supported by the National Science Foundation Living Stock Collections for Biological Research Program (#0951671 and #00017383). This work was supported by National Science Foundation Grant MCB #1818383 to John R. Pringle and M.O., National Science Foundation CAREER #2337141 to M.O., and a Duke University Arts and Sciences Faculty Research Grant to M.O. D.S. was a recipient of the Duke Biological Sciences Undergraduate Research Fellowship. S-A.C. was a recipient of the Hung Taiwan-Duke University Fellowship. Support for E.C. was provided by the Mentorship Program at the North Carolina School of Science and Mathematics.

## Abbreviations

EGT: endosymbiotic gene transfer
RGT: reverse gene transfer
CHES: N-cyclohexyl-2-aminoethanesulfonic acid
CCM: carbon-concentration mechanism
cTP: chloroplast transit peptide
NG: NeonGreen
SC: Scarlet.

## Author Contributions

Conceived and Designed Experiments. S-A.C., P.H., V.B., A.K., and M.O.

Performed the experiments. D.S., S-A.C., and E.C.

Analyzed the Data. D.S., S-A.C., P.H., and M.O.

Drafted the Article. D.S., S-A.C., and M.O.

Prepared the Digital Images. D.S., S-A.C., and M.O.

---

**SUPPLEMENTAL FIGURE S1:**
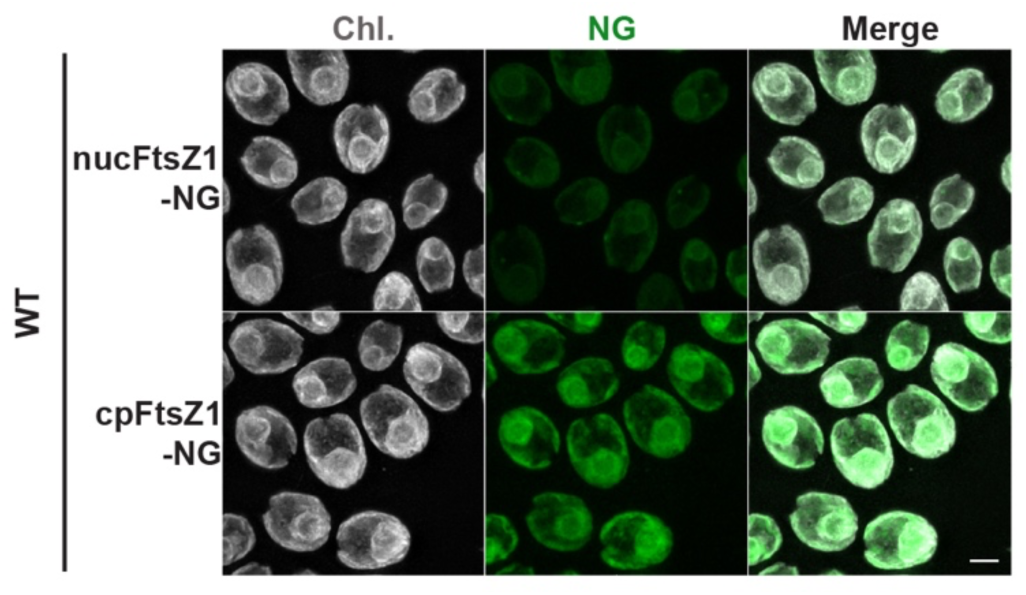
Time-course imaging of MOC399 (*nucFTSZ1-NG*) and DSC029 (*cpFTSZ1-NG*) during interphase of the cell cycle. The cells were synchronized on TAP agar. The cells were imaged at hour 5.5 of the 12L:12D cycle (mid-G1 phase, before endogenous *FTSZ1* is upregulated). Scale bar, 5 µm.

**SUPPLEMENTAL FIGURE S2:**
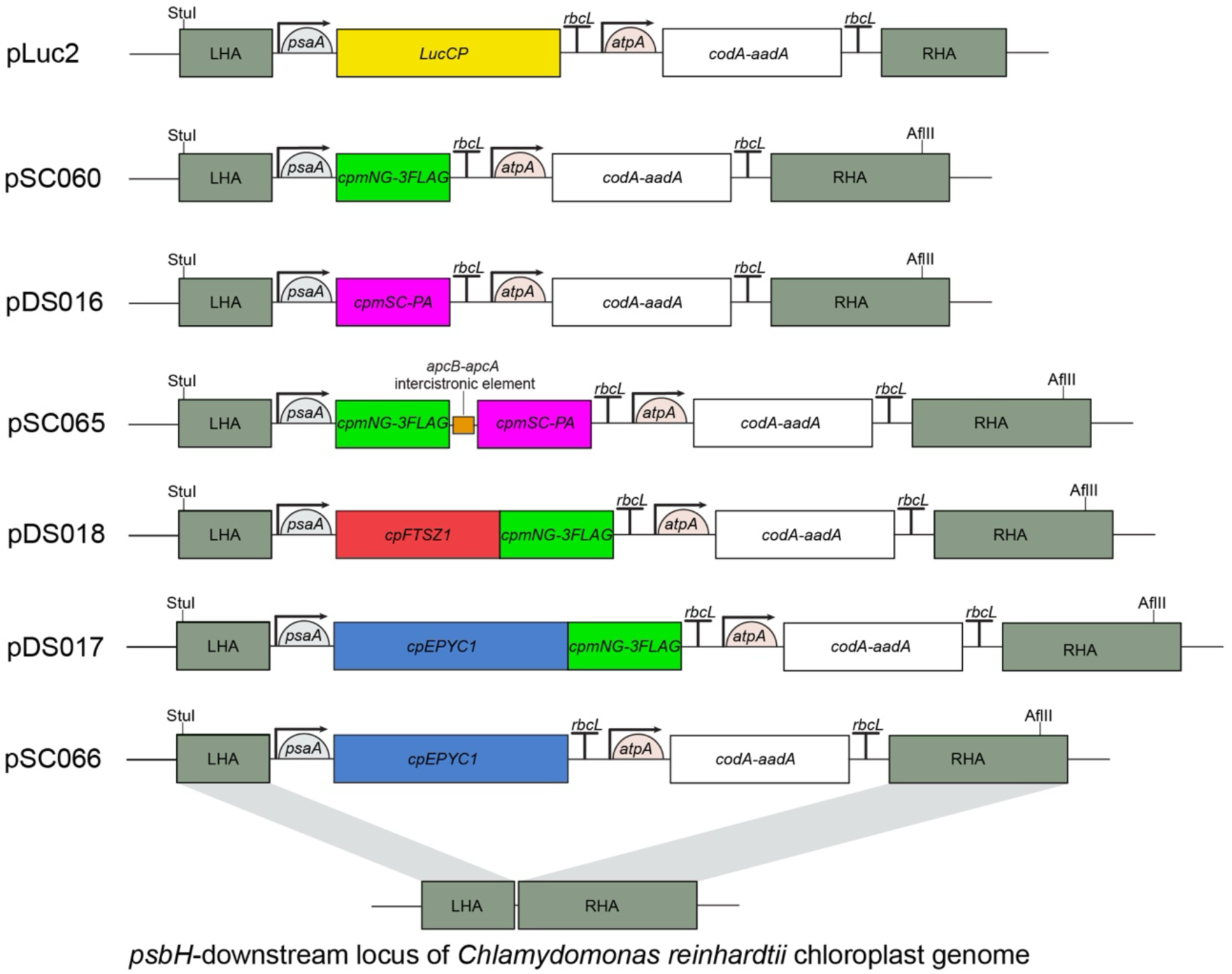
Schematic maps of chloroplast expression constructs. Each construct contains left and right homology arms (LHA and RHA, for homologous recombination at the *psbH*-downstream locus of the *Chlamydomonas reinhardtii* chloroplast genome. Indicated chloroplast codon-optimized transgenes are flanked by the *psaA* promoter/5′ UTR and the *rbcL* 3′ UTR/terminator. The *codA-aadA* selectable marker is flanked by the *atpA* promoter/5′ UTR and the *rbcL* 3′ UTR/terminator.

**SUPPLEMENTAL TABLE 1:**
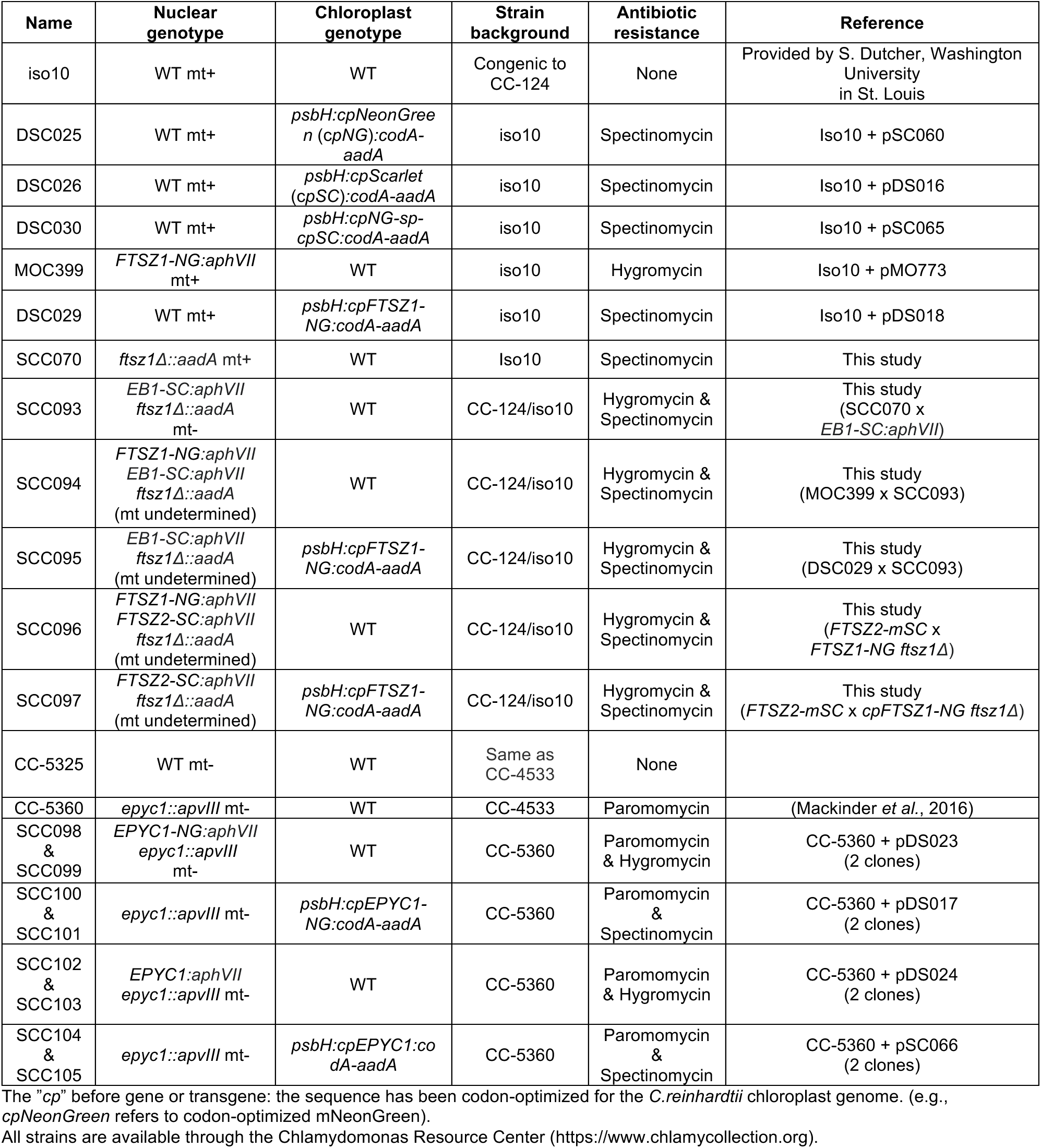
List of strains used in this study.

**SUPPLEMENTAL TABLE 2:** List of plasmids used in this study.

| Plasmid Name | Expression Cassette | Selection Marker<br>(*Chlamydomonas*) | Description |
| --- | --- | --- | --- |
| pALT007 |  | *codA-aadA* | The acceptor plasmid for generation of chloroplast expression plasmids targeted for homologous recombination at the *C. reinhardtii* chloroplast *psbH* locus. |
| pSC056 | *cpNG_v2-3FLAG* |  | Plasmid containing the codon-optimized *mNeonGreen* (*cpNG*) with *SapI* digestion sites at the two ends. Used to generate pSC060 through Start-Stop assembly. |
| pSC060 | *P<sub>psaA</sub>:cpNG_v2-3FLAG:rbcL-3'UTR* | *codA-aadA* | The expression plasmid of *cpNG* flanked by *psaA* promoter and *rbcL* 3'UTR; targeted for recombination at the *pshH* locus in chloroplast genome. |
| pSC057 | *cpSC_v2-PA* |  | Plasmid containing the codon-optimized *mScarlet* (*cpSC*) with *SapI* digestion sites at the two ends. Used to generate pDS016 through Start-Stop assembly. |
| pDS016 | *P<sub>psaA</sub>:cpSC_v2-PA: rbcL-3'UTR* | *codA-aadA* | The expression plasmid of *cpSC* targeted for recombination at the *pshH* locus. |
| pSC065 | *P<sub>psaA</sub>: cpNG_v2-3FLAG-spacer-cpSC_v2-PA: rbcL-3'UTR* | *codA-aadA* | The expression plasmid containing the bicistronic expression cassette of *cpNG* and *cpSC*; the two fluorophores are separated by a spacer in the *apcA/apcB* operon of *S. elongatus*. |
| pSC058 | *cpFTSZ1* |  | Plasmid containing the codon-optimized *cpFTSZ1* with *SapI* digestion sites at the two ends. |
| pSC059 | *cpEPYC1* |  | Plasmid containing the codon-optimized *cpEPYC1* with *SapI* digestion sites at the two ends. |
| pSC055 | *cpFTSZ1-NG_v2-3FLAG* |  | Plasmid containing the *cpFTSZ1-NG* with *SapI* digestion sites at the two ends. Used to generate pDS018 through 4-way Start-Stop assembly. |
| pDS018 | *P<sub>psaA</sub>:cpFTSZ1-NG_v2-3FLAG: rbcL-3'UTR* | *codA-aadA* | The expression plasmid of transgene *cpFTSZ1-NG*. |
| pDS015 | *cpEPYC1-NG_v2-3FLAG* |  | Plasmid containing the *cpEPYC1-NG* with *SapI* digestion sites at the two ends. Used to generate pDS018 through 4-way Start-Stop assembly. |
| pDS017 | *P<sub>psaA</sub>:cpEPYC1-NG_v2-3FLAG: rbcL-3'UTR* | *codA-aadA* | The expression plasmid of transgene *cpEPYC1-NG*. |
| pSC066 | *P<sub>psaA</sub>:cpEPYC1:rbcL-3'UTR* | *codA-aadA* | The expression plasmid of transgene *cpEPYC1*. |
| pDS023 | *P<sub>PsaD</sub>:EPYC1-NG_v2-3FLAG* | *HygR* | Nuclear expression plasmid of *EPYC1-NG*. |
| pDS024 | *P<sub>PsaD</sub>:EPYC1* | *HygR* | Nuclear expression plasmid of *EPYC1*. |

All plasmids are available through the Chlamydomonas Resource Center (https://www.chlamycollection.org).

